# Enhancement of synaptic AMPA receptors depends mutually on Src and PSD-95

**DOI:** 10.1101/2020.10.02.323568

**Authors:** Xiaojie Huang, Juliane M. Krüger, Anna Beroun, Weifeng Xu, Yan Dong, Oliver M. Schlüter

## Abstract

Synaptic incorporation and removal of AMPA receptors is highly regulated to modulate the strength of synaptic transmission for long-term synaptic plasticity during brain development and associative learning. PSD-93α2 and PSD-95α, two paralogs of the DLG-MAGUK protein family of signaling scaffolds govern the synaptic incorporation and stabilization of AMPA receptors opposingly, with PSD-95α promoting and PSD-93α2 inhibiting it. The associated signaling mechanisms that control the synaptic incorporation and stabilization remain elusive. Here, we used domain swapping between the antagonizing signaling scaffolds to identify the protein motifs responsible for enhancing synaptic AMPA receptors and the associated signaling protein. We narrowed down multiple motifs in the N-terminal domain that are principally responsible for governing the enhancement by Src. Specific activation and inhibiting peptides revealed continuous activity of Src. Together, the results depict a mutual dependence of Src and PSD-95α in enhancing and maintaining synaptic AMPA receptors.

## Introduction

Synaptic incorporation and removal of AMPA receptors is highly regulated to modulate the strength of synaptic transmission. This plasticity is thought to alter and shape excitatory network configurations to store information as patterns of synaptic weights and is recalled by activation of specific neuronal ensembles (Basu and Siegelbaum, 2015; Nicolelis et al., 1997). Developmental plasticity as well as Hebbian plasticity are forms of NMDA receptor-dependent long-term synaptic plasticity and are primarily expressed as changes in synaptic AMPA receptor numbers (Huganir and Nicoll, 2013; Kessels and Malinow, 2009; Malenka and Bear, 2004; Morris, 2013; Neves et al., 2008; Tsien et al., 1996; Xu et al., 2020). Despite its fundamental importance, the understanding of the signaling mechanisms that regulate AMPA receptor incorporation and removal, remain controversial and incompletely understood. While early work focused on changes at AMPA receptors themselves, later studies revealed that posttranslational modifications on the receptors appear rather permissive than instructive and triggered the concept that signaling events through signaling scaffolds in the postsynaptic density (PSD) determine the AMPA receptor content (Granger et al., 2013; Hayashi et al., 2000; Lee et al., 2003; Shi et al., 2001; Xu et al., 2008; Zamanillo et al., 1999).

The disc large (DLG)-membrane associated guanylate kinase (MAGUK) family of proteins, PSD-93/Chapsyn-110, PSD-95/synapse associated protein (SAP)90, SAP97 and SAP102, are signaling scaffolds of the PSD, which regulate AMPA receptor trafficking in the synapse during basal synaptic transmission and long-term synaptic plasticity (Béïque et al., 2006; Carlisle et al., 2008; Cuthbert et al., 2007; Ehrlich et al., 2007; Elias et al., 2006; Liu et al., 2017; Migaud et al., 1998; Nakagawa et al., 2004; Schlüter et al., 2006; Steiner et al., 2008; Xu et al., 2008). The principal isoforms of PSD-93 and PSD-95 (α-isoforms) share high sequence similarity with a palmitoylated N-terminal domain, three PDZ domains, an SH3 domain and a C-terminal guanylate kinase (GK) domain (Brenman et al., 1996; Cho et al., 1992; Kim et al., 1996; Krüger et al., 2013; Parker et al., 2004). In contrast, the principal isoform of SAP97 (β-isoforms) has an N-terminal L27 domain, while SAP102 has a unique N-terminal domain with both paralogs sharing the rest of the overall domain structure (Müller et al., 1996; Müller et al., 1995; Schlüter et al., 2006). The palmitoylated α-isoforms of PSD-95 and SAP97 generally enhance synaptic incorporation of AMPA receptors (Béïque and Andrade, 2003; Elias et al., 2006; Nakagawa et al., 2004; Schlüter et al., 2006; Schnell et al., 2002b), but PSD-95α also regulates AMPA receptor removal during NMDA receptor-dependent long-term synaptic depression (Béïque and Andrade, 2003; Xu et al., 2008). Thus, the same signaling scaffold is involved in both, stabilization of synaptic AMPA receptors and their removal, indicating multiple and opposing functionalities harbored in one protein.

In the developing brain, the α-isoforms of PSD-95 and PSD-93 regulate synaptic AMPA receptor incorporation opposingly, with PSD-95α promoting and PSD-93α2 inhibiting the incorporation (Béïque et al., 2006; Carlisle et al., 2008; Favaro et al., 2018; Krüger et al., 2013). The molecular underpinnings of the opposing function are unknown and the distinct signaling mechanisms of PSD-95α to regulate synaptic AMPA receptor incorporation remain elusive.

Members of the sarcoma (Src) family protein kinases were found to regulate long-term synaptic plasticity and the function of DLG-MAGUKs (Grant et al., 1992; Nada et al., 2003; Sato et al., 2008). Src family protein kinases reside in different activation states, dependent on ligand binding to their regulatory domains and their phosphorylation states, thus operate as graded switches (Bradshaw, 2010). Long-term synaptic potentiation (LTP) induction activates Src, which enhances NMDA receptor activities and facilitates LTP (Lu et al., 1998). PSD-95 binds to Src, but whether this binding inhibits or activates Src-mediated NMDA receptor phosphorylation is controversial (Kalia et al., 2006b; Kalia and Salter, 2003; Zhang et al., 2008; Zhao et al., 2015). Besides its effects on NMDA receptors and LTP, it remains elusive whether Src regulates synaptic AMPA receptor incorporation independent of regulating NMDA receptor function.

Here, we used domain swapping between PSD-93α2 and PSD-95α or SAP97α to identify the protein motifs responsible for enhancing synaptic AMPA receptor-mediated transmission. We found that multiple motifs in the N-terminal domain of PSD-95α or SAP97α are principally responsible for governing the enhancement. These motifs are associated with Src activation, leading to a mutual dependence of Src and PSD-95α in enhancing and maintaining synaptic AMPA receptors.

## Material and Methods

### Virus vectors and neuronal transduction

All lentiviral constructs were made from the lentiviral transfer vector FUGW (Lois et al., 2002) and its variant FHUG+W, which additionally contains an RNAi expression cassette driven by an H1 promoter for knocking down PSD-95 (sh95RN) or Src (shSRf) (Schlüter et al., 2006). C-terminally eGFP-tagged PSD-93α2 and SAP97α over-expression and PSD-95-to-PSD-95α replacement constructs were described previously (Krüger et al., 2013; Schlüter et al., 2006).

Transduction of neurons in rat hippocampal slice cultures was achieved by injecting concentrated viral particles into the CA1 pyramidal cell layer on DIV 1 or DIV 2 using a Nanoject II device (Drummond Scientific) (Krüger et al., 2013).

### Construction of lentiviral vectors

For the analysis of the differences in the N-termini of PSD-93α and PSD-95α, exon 1 of PSD-93α2 was replaced by exon 1 of PSD-95α in the lentiviral over-expression constructs, and vice versa, leading to the constructs PSD-93_95_1-9_ and PSD-95_93_1-20_, respectively. To study the differences in the N-terminus upstream of PDZ-1, the N-terminus of PSD-93α2 (amino acids aa1-104) was replaced by the N-terminus of PSD-95*α* (aa 1-64) or N-terminus of SAP97α (aa1-94) and vice versa, leading to the constructs PSD-93_95_1-64_ or PSD-93_97_1-94_ and PSD-95_93_1-104_. Exon 2 of SAP97α (aa11-65) was replaced with exon 2 of PSD-93α2 (aa22-75), and vice versa, leading to constructs SAP97_93/E2_22-75_ and PSD-93_97/E2_11-65_, respectively. The extended exon 3 of SAP97α (aa66-94) was replaced with that of PSD-93α2 (aa76-104), and vice versa, leading to the constructs SAP97_93/E3_76-104_ and PSD-93_97/E3_66-94_, respectively. To substitute F103 with Tyr in PSD-93α2, we used standard site directed mutagenesis to change the codon TTT to TAC to generate the over-expression construct PSD-93FY. In the PSD-95 wild-type replacement construct, we used site directed mutagenesis to introduce the Y63F mutation or the G59I and Y63F double mutation to generate the replacement constructs PSD-95YF and PSD-95IF, respectively. The identity of each construct was confirmed by DNA sequencing.

### Hippocampal slice culture

All procedures were performed by following the procedures approved by the animal care and use committees and governmental agencies of the listed institutions. Organotypic hippocampal slices were prepared according to previously published procedures (Krüger et al., 2013; Schlüter et al., 2006). In short, P7-P9 Wistar rats of either sex were anesthetized with isofluorane, and the hippocampi were dissected in ice-cold sucrose cutting buffer (in mM as follows: 204 sucrose, 26 NaHCO_3_, 10 glucose, 2.5 KCl, 1 NaH_2_PO4, 4 MgSO_4_, 1 CaCl_2_, 4 ascorbic acid). Using a guillotine with a wire frame, 300-µm-thick slices were cut and transferred into artificial CSF (ACSF) (containing in mM) as follows: 119 NaCl, 26 NaHCO_3_, 20 glucose, 4 MgSO_4_, 2.5 KCl, 1 NaH_2_PO_4_, 4 CaCl_2_, for recovery. The slices were washed in BME (Biochrom) and plated onto MilliCell (Millipore) culture plate inserts in culture dishes containing BME-based culture medium supplemented with 20% heat-inactivated horse serum (Biochrom). The slices were incubated at 37°C in a 5% CO_2_ environment. On DIV 1, the slices were transferred to 34°C, and the culture medium was refreshed.

Transduction of neurons in hippocampal slice cultures was achieved by injecting concentrated viral particles into the CA1 pyramidal cell layer on DIV 1 or DIV 2 using a Nanoject II device (Drummond Scientific).

### Acute hippocampus slice preparation

300um thick coronal acute brain slices were prepared from P18-20 mouse using a virbrating microtome (Leica VT1200S) in ice cold cutting solution containing in mM (119 Choline Cl, 26 NaHCO_3_, 30 Glucose, 7 MgSO_4_, 2.5 KCl, 1 NaH_2_PO_4_, 1 CaCl_2_, 1 Kynurenic acid, 1.3 Na ascorbate, 3 Na pyruvate). Slices were recovered in oxygenated ACSF (119 NaCl, 26 NaHCO_3_, 20 Glucose, 1.3 MgSO_4_, 2.5 KCl, 1 NaH_2_PO_4_, 2.5 CaCl_2_) at 34°C for 30 minutes, and then were kept at room temperature until use.

### Electrophysiology

Slice culture recordings were performed 4-9 days after viral vector transduction as previously described (Krüger et al., 2013; Schlüter et al., 2006). A single slice was transferred into the recording chamber, which was constantly perfused with oxygenated ACSF (2-3 ml/min) containing in mM (119 NaCl, 26 NaHCO_3_, 20 Glucose, 4 MgSO_4_, 2.5 KCl, 1 NaH_2_PO_4_, 4 CaCl_2_). 1-5 µM 2-chloroadenosine (Biolog) was added to the ACSF to reduce polysynaptic activity. 50 µM picrotoxin (Ascent Scientific) was added to block inhibitory transmission and thus to isolate excitatory postsynaptic currents (EPSCs). Patch pipettes (open pipette resistance 3-6 MOhm) were filled with intracellular solution consisting of (in mM) 117.5 MeSO_3_H, 10 HEPES, 17.75 CsCl, 10 TEA-Cl, 0.25 EGTA, 10 Glucose, 2 MgCl_2_, 4 Na_2_ATP, 0.3 NaGTP (pH 7.3, 290 mOsm). For pairwise recording, an infected cell and its neighboring uninfected (control) cell were simultaneously recorded using whole-cell voltage-clamp configuration. EPSCs were simultaneously evoked in paired cells by a single bipolar stimulation electrode filled with ACSF. AMPAR EPSCs were recorded at −60 mV. For NMDAR EPSCs, EPSCs were recorded at +40 mV, and the amplitude of NMDAR EPSCs was assessed by the amplitude of EPSCs at 60 msec after the peak currents, a time point at which AMPAR EPSCs are mostly absent.

Acute slice recordings were performed with similar ACSF except using 1.3 MgSO_4_ and 2.5 CaCl_2_. Cs-gluconate based intracellular solution containing in mM (130 Cs gluconate, 20 HEPES, 10 EGTA, 3 TEA-Cl, 4 QX314-Cl, 4 MgATP, 0.3 Na_2_GTP) were used.

Signals were filtered at 4 kHz and digitalized at 10 kHz using an ITC-18 acquisition board (HEKA). The responses were monitored online and analyzed using custom-written acquisition and analysis software implemented in IGOR Pro (Wavemetrics). For each cell pair, a minimum of 40 sweeps was collected and averaged. The stimulation strength was adjusted such that the amplitude of AMPAR EPSCs in control cells was ∼100 pA. A 5 mV hyperpolarizing step was used to monitor the input and series resistance over the recording.

The statistical analysis of electrophysiological data was done using the software Excel (Microsoft), Igor (Wavemetrics) or Prism (GraphPad). Comparison between pairs of control and infected cell responses were analyzed using the two-tailed paired T test. For none paired measures, two-tailed T test with unequal variance was used. Comparison between more than two constructs was performed using the ANOVA one factor test followed by Dunnett posthoc test. For Src activation peptide experiment, mixed model two-way ANOVA was used to check the difference between control and experiment group with the peptide. All data are presented as mean ± standard error of the mean (SEM), which is shown as error bars in the graphs.

## Results

### N-terminal domains of PSD-95α and PSD-93α2 transfer their differential roles in regulating AMPA receptor function

Unlike PSD-95α and SAP-97α, over-expression of the two N-terminal α-isoforms of PSD-93 does not enhance AMPA receptor-mediated (R) EPSCs (Favaro et al., 2018; Krüger et al., 2013; Liu et al., 2014; Nakagawa et al., 2004; Schnell et al., 2002b). Conversely, gene deletion or knock-down of *Dlg4* (PSD-95), but not *Dlg2* (PSD-93) reduces AMPAR EPSCs (Béïque et al., 2006; Carlisle et al., 2008; Favaro et al., 2018; Krüger et al., 2013; Liu et al., 2018; Nakagawa et al., 2004). The N-terminus of PSD-95 that contained the first two PDZ domains enhances AMPAR EPSCs if over-expressed (Schnell et al., 2002b; Xu et al., 2008). To explore the structural basis for this functional difference between the N-terminus of PSD-93α2 and PSD-95α or SAP97α, we compared the amino acid sequences and protein structure of the three DLG-MAGUKs. The first two PDZ domains are highly conserved. At the N-terminus, the highest variability is in the exon 2, which in *Dlg4* only extends shortly over the conserved motif in α and β-DLG-MAGUKs that translates to KYRYQDED (Fig. 1A). Exon 2 in *Dlg2* and *Dlg1* (SAP97) is twice as long as in *Dlg4*. However, the length per se does not determine the functional difference, because over-expression of SAP97α enhances AMPAR function (Schlüter et al., 2006; Schnell et al., 2002b), whereas PSD-93αs do not (Favaro et al., 2018; Krüger et al., 2013). However, one study implicates functional redundancy of PSD-95 and −93 in supporting synaptic AMPAR EPSCs (Elias et al., 2006).

**Figure 1:**
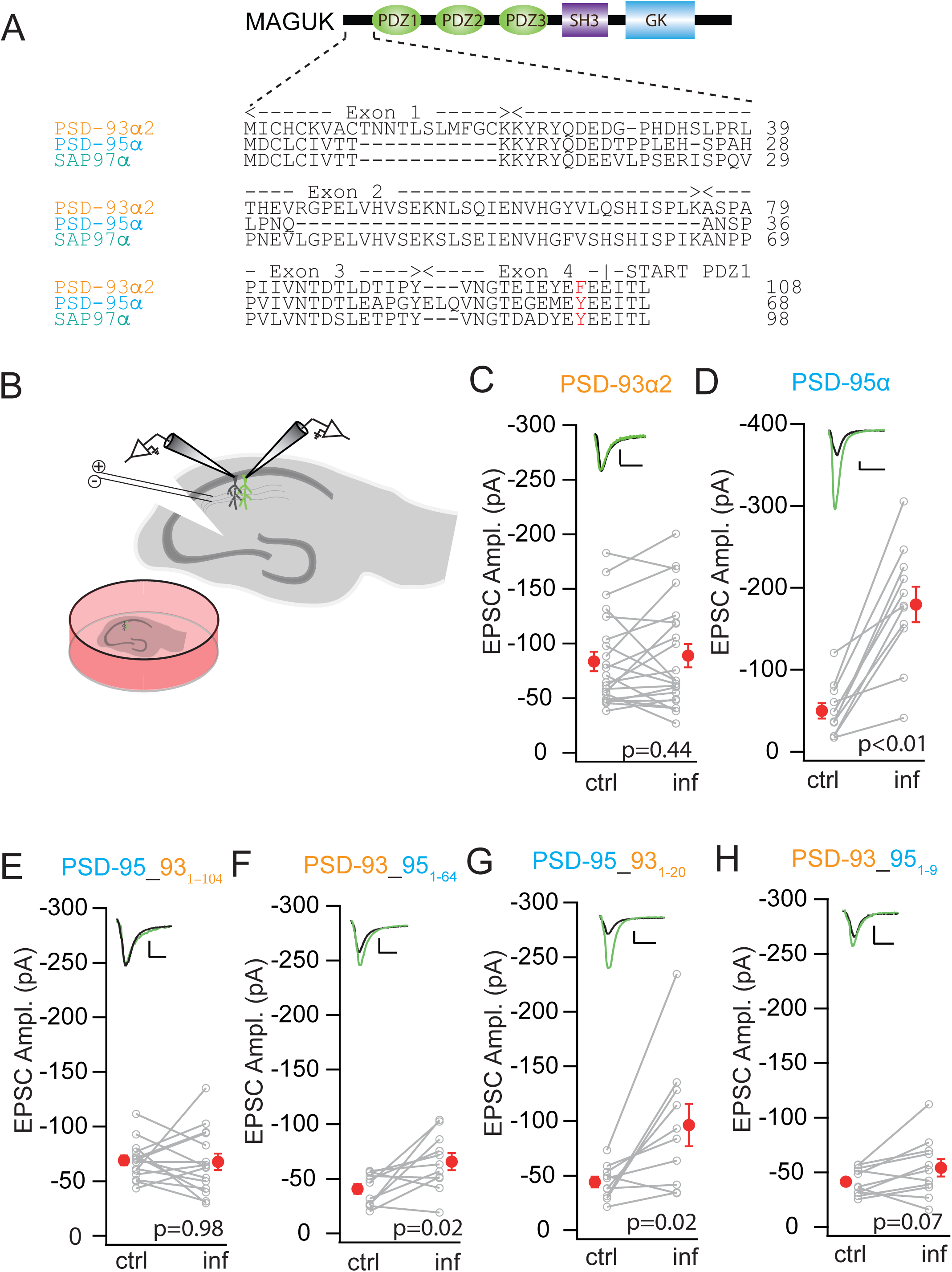
N-terminal domains of PSD-93α2 and PSD-95α mediate specific function on AMPAR EPSCs. (A) Alignment of N-terminal amino acid sequence of PSD-93α2, PSD-95α, and SAP97α. Exon borders are indicated, and a conserved Y in PSD-95α and SAP97α, while a F in PSD-93α2 highlighted in red. Domain structure of the DLG-MAGUKs is depicted on top. (B) Schematic drawing of the recording configuration of organotypic slices from rat hippocampus. (C-H) Amplitude of AMPAR EPSCs of CA1 pyramidal neurons expressing eGFP-tagged (inf) (C) PSD-93α2, (D) PSD-95α, (E) PSD-95 with PSD-93α2 N-terminal domain, (F) PSD-93 with PSD-95α N-terminal domain (G) PSD-95 with PSD-93α2 palmitoylation motif, and (H) PSD-93 with PSD-95α palmitoylation motif as lentiviral constructs are plotted against those of simultaneously recorded uninfected neighboring control neurons (ctrl). In this and all subsequent panels: gray symbols linked by a line represent single pairs of recordings; red symbols depict mean ± SEM, p values calculated by paired t-test. Insets in each panel depict sample traces of control (black) and transduced neurons (green), Scale bar: 25 pA, 20 ms.).

To establish the baseline observation to compare and contrast domain swapping mutants, we over-expressed PSD-93α2 and PSD-95α, and measured AMPAR EPSCs with the dual whole cell patch-clamp procedure, simultaneously recording from a transduced CA1 pyramidal neuron and a neighboring reference neuron in rat organotypic slice cultures (Fig. 1B). This recording configuration allows the quantitative comparison of AMPAR EPSCs between the two conditions (Hayashi et al., 2000; Schlüter et al., 2006). Consistent with our previous results in mouse visual cortex and rat organotypic hippocampal slice cultures, over-expression of PSD-93α2 did not alter AMPAR EPSCs (n = 22; control, −83.47 ± 8.93 pA; infected, −88.76 ± 10.66 pA, p = 0.44; Fig. 1C) (Krüger et al., 2013). In contrast, over-expression of PSD-95α enhanced AMPAR EPSCs (n = 11; control, −49.88 ± 9.61 pA; infected, −179.53 ± 21.76 pA, p < 0.01; Fig. 1D).

As a construct of PSD-95 with the N-terminus and the first two PDZ domains enhanced AMPAR EPSCs (Schnell et al., 2002b; Xu et al., 2008), we over-expressed a counterpart construct of PSD-93*α*2. However, the N-terminal PSD-93 construct had no such effect, indicating that the N-terminus of PSD-93 lacks motifs for enhancing AMPARs (n = 11; control, −60.78 ± 4.61 pA; infected, −48.66 ± 7.41 pA, p = 0.14). To locate and characterize these critical regulatory motifs, we constructed chimeras of PSD-95α and PSD-93α2. If we swapped the N-terminal domain up to PDZ-1 of PSD-93α2 onto PSD-95, the enhancing effect of PSD-95 on AMPARs was abolished (n = 15; control, −68.98 ± 4.49 pA; infected, −67.70 ± 7.60 pA, p = 0.88; Fig. 1E). This result echoed the result of the N-terminal PSD-93 construct that the AMPAR enhancing motifs in the N-terminal domain before the first PDZ domain are absent in PSD-93*α*2. We then did the reverse, swapping the N-terminus of PSD-95*α* onto PSD-93. When the full N-terminal domain (up to PDZ-1) was swapped, expression of this construct increased the peak amplitude of AMPAR EPSCs (n = 11; control, −40.73 ± 4.41 pA; infected, −65.74 ± 7.76 pA, p < 0.05; Fig. 1F). Together, these results indicated that the N-terminal region before the first PDZ domain in PSD-93*α*2 does not contain the functionality of enhancing AMPAR EPSCs, whereas the N-terminal region of PSD-95*α* does.

To further narrow down the location, we swapped the first 20 amino acids of the N-terminus of PSD-93α2 onto PSD-95. This short sequence codes for the palmitoylation signal before the conserved KYRYQDED motif. Expression of this construct increased AMPAR EPSCs at a regulatory magnitude similar to PSD-95*α* (n = 10; control, −44.04 ± 4.92 pA; infected, −96.32 ± 19.41 pA, p = 0.02; Fig. 1G). Conversely, swapping the very first 9 amino acids before the KYRYQDED motif from PSD-95 onto PSD-93 did not enhance AMPAR EPSCs (n = 11; control, - 41.52 ± 3.64 pA; infected, 54.02 ± 8.11 pA, p = 0.07; Fig. 1H), indicating that the very N-terminus has little contribution to the functional difference between PSD-95α and PSD-93α2. Together, the above results indicated that one or more motifs between the conserved KYRYQDED and PDZ-1 were lacking in PSD-93, and these motifs were required for the enhancing effect of DLG-MAGUK α-isoforms on AMPARs.

### Motifs in consecutive exons coding the N-terminal domain contributed to the AMPAR-enhancing effect of DLG-MAGUK α-isoforms

Compared to PSD-95, SAP97 is more similar to PSD-93 in their N-terminal domains (Fig. 1A). Given over-expression of SAP97α upregulates AMPAR EPSCs, whereas over-expression of PSD-93α2 does not (Krüger et al., 2013; Schlüter et al., 2006), we reasoned that by swapping N-terminal exons between *Dlg2* and *Dlg1*, we might further narrow down the critical motifs governing the α-isoform-mediated enhancement of AMPARs. We first confirmed the enhancing effect of SAP97α (Schlüter et al., 2006). When SAP97α was over-expressed, AMPAR EPSCs were increased (n = 23; control, −66.96 ± 4.70 pA; infected, - 140.69 ± 10.99 pA, p < 0.01; Fig. 2B, I). The AMPAR-enhancing effect of SAP97α was transferred onto PSD-93α2 by swapping the SAP97α N-terminal domain onto PSD-93 (n = 18; control, −34.99 ± 4.07 pA; infected, −84.16 ± 11.53 pA, p < 0.01; Fig. 2C, F), echoing the result of the PSD-95/PSD-93 N-terminal swap and further supporting the N-terminus as the principal location of the enhancing effect.

**Figure 2:**
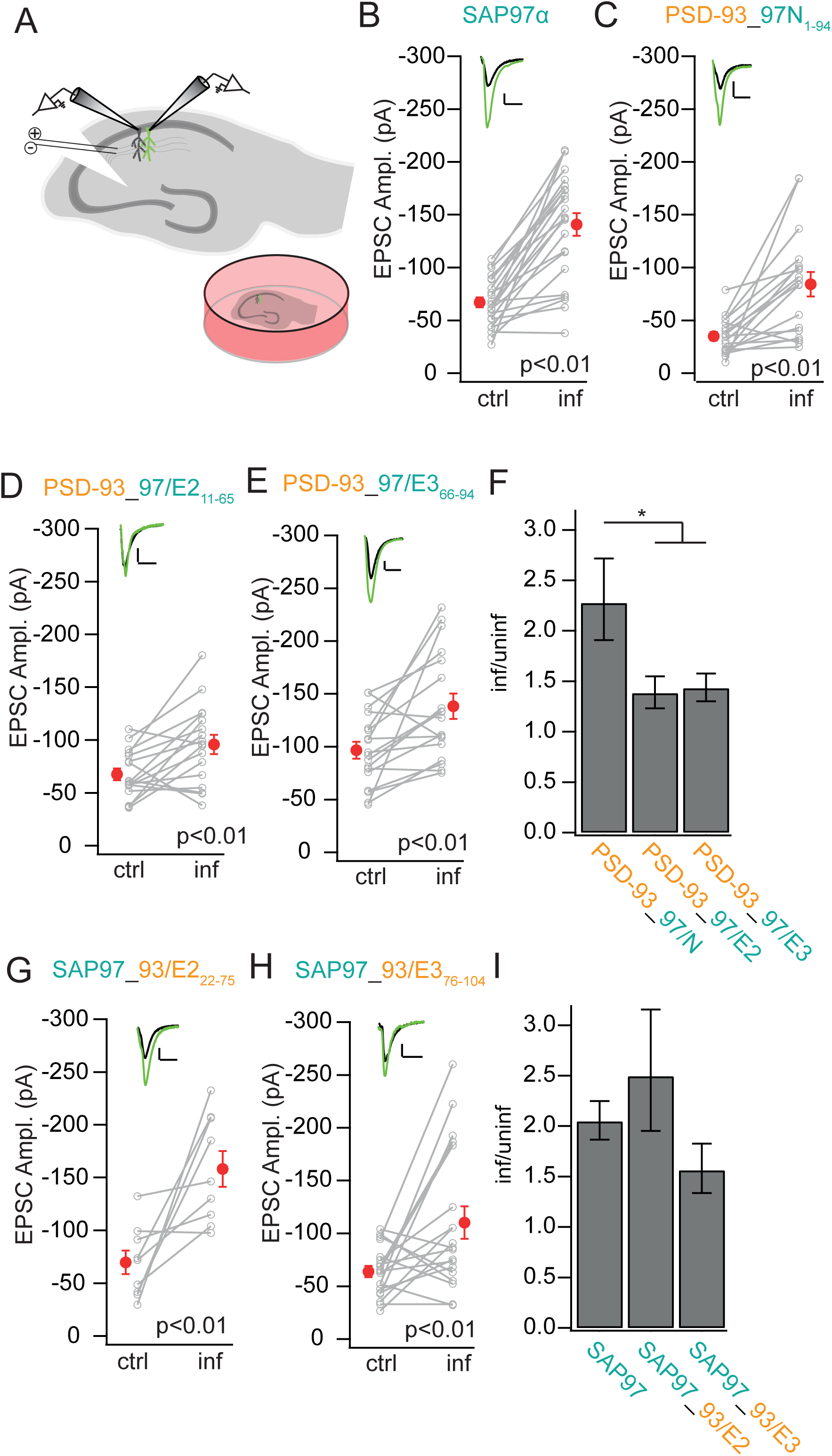
Multiple motifs in N-terminal domain contribute to the α-type enhancement. (A) Schematic drawing of the recording configuration of organotypic slices from rat hippocampus. (B-E,G,H) Amplitude of AMPAR EPSCs of CA1 pyramidal neurons expressing eGFP-tagged (B) SAP97α, (C) PSD-93 with SAP97α N-terminal domain (D) PSD-93 with SAP97α exon 2, (E) PSD-93 with SAP97α exon 3. (G) SAP97 with PSD-93α2 exon 2, and (H) SAP97 with PSD-93α2 exon 3, as lentiviral constructs are plotted against those of simultaneously recorded uninfected neighboring neurons in hippocampal slice cultures. (F, I) Summary graph of the comparison between indicated experimental groups of AMPAR EPSC amplitude ratios between transduced (inf) and control neuron (uninf). ANOVA test: * p < 0.05.

The N-terminal swaps so far narrowed the enhancing effect down to exons 2, 3 and parts of exon 4 up to the coding region of PDZ-1. In a next step, we split the N-terminus at the exon 2 to exon 3 boundary to either swap exon 2 or exon 3 with parts of exon 4 (referred to as extended exon 3 in the following). The PSD-93 construct with exon 2 of *Dlg1* (PSD-93_97/E2_11-65_) increased the peak amplitude of AMPAR EPSCs, but with a comparatively low magnitude (n = 17; control, −67.46 ± 5.46 pA; infected, −95.67 ± 9.17 pA, p < 0.01; Fig. 2D). The PSD-93 construct with extended exon 3 of *Dlg1* (PSD-93_97/E3_66-94_) increased the peak amplitude of AMPAR EPSCs similar to the exon 2 swap (n = 18; control, −96.57 ± 7.91 pA; infected, −138.24 ± 11.95 pA, p < 0.01; Fig. 2E).

When comparing the magnitudes of effect of the swaps of the partial N-terminal motifs, coded by exon 2 (PSD-93_97/E2_11-65_) or extended exon 3 (PSD-93_97/E3_66-94_) with the full N-terminal swap (PSD-93_97/N), the magnitude of enhancement of the partial swaps was smaller (PSD-93_97/N, 2.27 ± {+ 0.44, - 0.37}; PSD-93_97/E2, 1.38 ± {+0.17, −0.15}; PSD-93_97/E3, 1.43 ± {+0.14, −0.13}; F_(2,50)_ = 4.29, p = 0.02; PSD-93_97/N vs. PSD-93_97/E2, p = 0.01; PSD-93_97/N vs. PSD-93_97/E3, p = 0.02; Fig. 2F), indicating that both exon 2 and extended exon 3 of *Dlg1* contributed to the enhancement additively.

In the converse experiment, the SAP97 construct with *Dlg2* exon 2 (SAP97_93/E2_22-75_) increased the peak amplitude of AMPAR EPSCs (n = 9; control, −69.79 ± 11.21 pA; infected, −158.23 ± 16.83 pA, p < 0.01; Fig. 2G). Similarly, the SAP97 construct with *Dlg2* extended exon 3 (SAP97_93/E3_76-104_) increased the peak amplitude of AMPAR EPSCs (n = 19; control, −63.81 ± 5.36 pA; infected, −110.28 ± 15.29 pA, p < 0.01; Fig. 2H). Comparison of the magnitude of effect of SAP97α with the SAP97 constructs with *Dlg2* exon 2 (SAP97_93/E2_22-75_) or extended exon 3 (SAP97_93/E3_76-104_) revealed no significant difference and only swapping the *Dlg2* extended exon 3 exhibited a trend of smaller enhancement (SAP97, 2.04 ± {+0.20, −0.18}; SAP97_93/E2, 2.40 ± {+0.67, - 0.52}; SAP97_93/E3, 1.56 ± {+0.26, −0.22}; F_(2,48)_ = 1.91, p = 0.16; Fig. 2I). Thus, SAP97 with either its exon 2 or extended exon 3 enhanced AMPAR EPSCs fully, while PSD-93 required both exon 2 and extended exon 3 of either PSD-95 or SAP97 for the enhancing effect (Fig. 2). We interpret this different outcome by potential additional motifs in the C-terminal domain that might cooperate with motifs in the N-terminal domain to either enhance with the SAP97 C-terminal domain or inhibit with the PSD-93 C-terminal domain the enhancing function. Because the AMPAR-enhancing motif was transferred by the N-terminus, which is not part of the palmitoylation motif, the AMPAR-enhancing motif should be either one that spans over exon 2 and 3 or a complex comprising of two relatively independent parts, one in exon 2 and the other in the extended exon 3.

### A single amino acid differentiated PSD-93 from PSD-95 and SAP97 in regulating AMPARs

Our results so far narrow down that exon 2 and extended exon 3 in *Dlg4* and *Dlg1* code for the likely motifs that entail the enhancing effect on synaptic AMPAR function. Thus, the critical amino acid sequence or even single amino acids that mediates the AMPAR-enhancing effect should exist in exon 2 and/or extended 3 in the α-isoform of PSD-95 and SAP97, but not in PSD-93α2. To identify a motif, we searched for conserved motifs and amino acids that are shared between PSD-95α and SAP97α and different in PSD-93α2. Exon 2 was very similar between SAP97 and PSD-93, whereas large parts were absent in PSD-95 (Fig. 1A). However, before the PDZ-1 domain in extended exon 3, there is a conserved tyrosine in PSD-95 (Y63) and SAP97 (Y93), whereas in PSD-93 this amino acid is a phenylalanine (F103). We created point mutations in PSD-93α2 and PSD-95α to test the role of this amino acid in α-isoform-mediated enhancement of synaptic AMPA receptor function. Because these mutations may not be dominant and can be masked by multimerization with endogenous PSD-95, we used the molecular replacement strategy to overcome this potential limitation (Xu et al., 2008). We first validated this strategy. Replacing the endogenous PSD-95 by recombinant PSD-95α, which resulted in over-expression of functional PSD-95, increased the peak amplitude of AMPAR EPSCs (n = 26; control, −60.43 ± 6.30 pA; infected, −202.24. ± 21.87 pA, p < 0.01; Fig. 3B, E). These results confirmed that the molecular replacement strategy was effective as we used previously (Schlüter et al., 2006; Xu et al., 2008). Replacing endogenous PSD-95 with PSD-95α containing the Y63F mutation reduced the magnitude of PSD-95α-mediated enhancement of AMPAR EPSCs. However, the molecular replacement with the Y63F mutant still enhanced AMPAR EPSCs (n = 21; control, −71.70 ± 5.68 pA; infected, −147.30 ± 15.53 pA, p < 0.01; Fig. 3C, E). In this sequence stretch of the extended exon 3, another difference between PSD-95 and PSD-93 exists, isoleucine (I99) in PSD-93 but glycine (G59) in PSD-95. SAP97 has an alanine at this position, which is not as bulky and hydrophobic as isoleucine, and thus might be more similar to the glycine of PSD-95. We created a double mutant of PSD-95α with G59I and Y63F to test whether the isoleucine has an additional effect on the α-isoforms. Similar to the Y63F single mutant, PSD-95 replacement with the double mutant enhanced AMPAR EPSCs (n = 16; control, −80.55 ± 7.82 pA; infected, −164.97 ± 17.93 pA, p < 0.01; Fig. 3D, E). The effect of PSD-95α WT replacement was bigger than either PSD-95α with the Y63F single mutation or with the G59I Y63F double mutation, and the mutant constructs did not differ from one another (PSD-95α, 3.30 ± {+0.61, −0.51}; PSD-95YF, 1.97 ± {+0.30, −0.26}; PSD-95IF, 2.03 ± {+0.31, −0.27}; F_(2,60)_ = 3.61, p = 0.03; PSD-95α vs. PSD-95YF, p = 0.02; PSD-95α vs. PSD-95IF, p = 0.04; Fig. 3E). Thus, the G59 or I99 appear irrelevant for the enhancing effect. However, Y63 is critical, but other sites in the α-isoforms of DLG-MAGUKs also contribute to the enhancement of synaptic AMPAR function.

**Figure 3:**
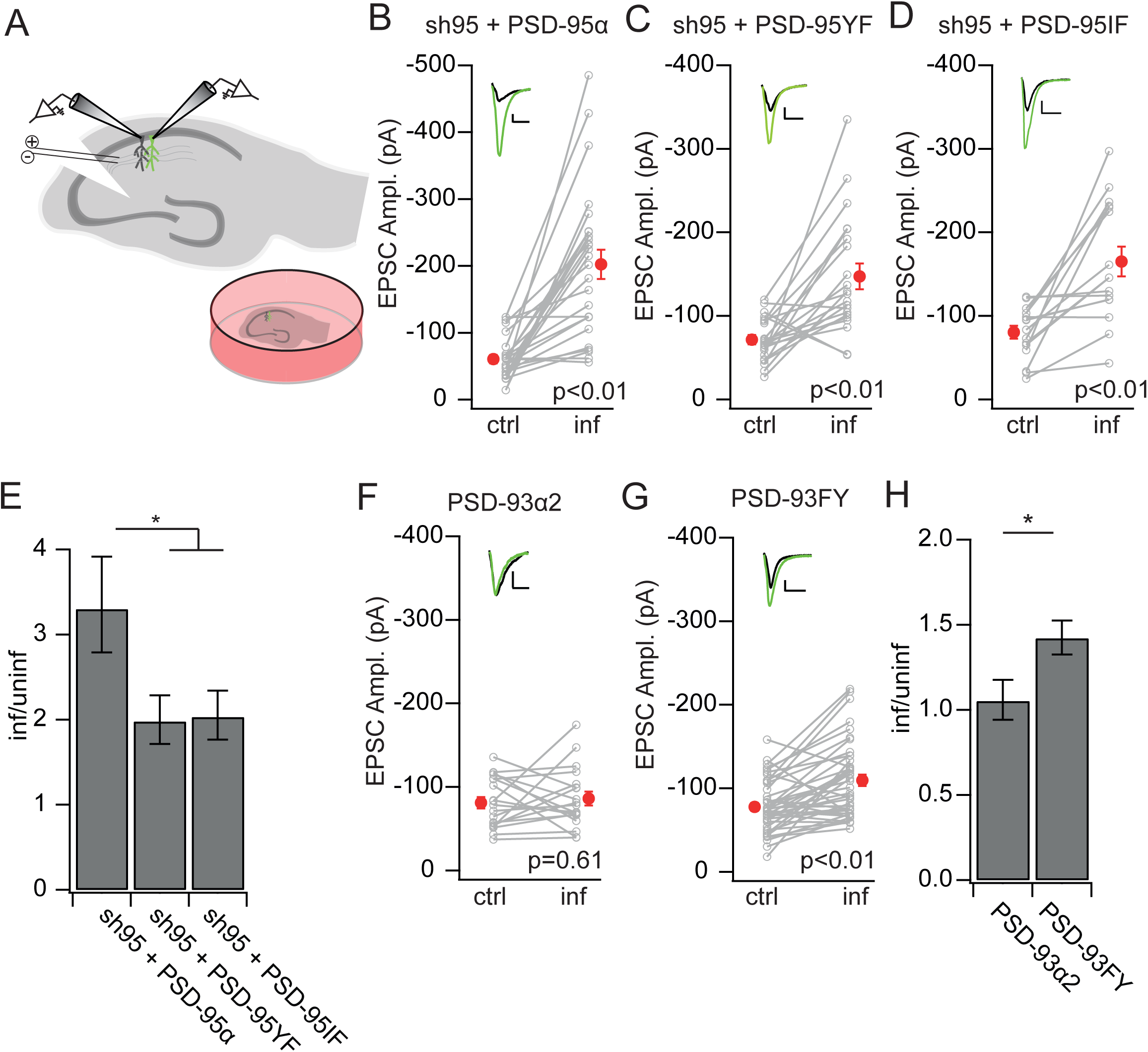
A single amino acid substitution of phenylalanine to tyrosine contributes to the α-type enhancement. (A) Schematic drawing of the recording configuration of organotypic slices from rat hippocampus. (B-D, F, G) Amplitude of AMPAR EPSCs of CA1 pyramidal neurons expressing eGFP-tagged (B) PSD-95α wild-type replacement, (C) PSD-95 Y63F replacement, (D) PSD-95 G59I Y63F replacement, (F) PSD-93α2 over-expression, and (G) PSD-93 F103Y over-expression as lentiviral constructs are plotted against those of simultaneously recorded uninfected neighboring neurons (ctrl). (E, H) Summary graph of the comparison between indicated experimental groups of AMPAR EPSC amplitude ratios between transduced (inf) and control neuron (uninf). ANOVA or T-test: * p < 0.05.

We then performed the reverse experiment. Over-expression of PSD-93α2 alone as interleaved recordings in this set of experiments had no significant effect on AMPAR EPSCs (n = 19; control, −81.01 ± 6.88 pA; infected, −86.06 ± 8.19 pA, p = 0.68; Fig. 3F). We created a mutant PSD-93α2 in which the F103 was substituted by tyrosine. Whereas over-expression of wild-type PSD-93α2 did not affect AMPAR EPSCs, over-expression of PSD-93α2 F103Y increased the peak amplitude of AMPAR EPSCs (n = 42; control, −77.76± 4.81 pA; infected, −109.58± 6.90 pA, p < 0.01; Fig. 3G), and its effect was different from over-expression of PSD-93α2 (PSD-93α, 1.05 ± {+0.12, −0.11}; PSD-93FY, 1.42 ± {+0.10, −0.09}; p = 0.03; Fig. 3H). These results again indicate the critical role of Y63 in the α-isoforms of DLG-MAGUKs to mediate the enhancement of synaptic AMPA receptor function, and again indicated that additional unidentified sequences/motifs, possibly in exon 2, are also involved in the regulation of AMPARs.

### SFK activating peptide required PSD-95 to enhance synaptic AMPA receptors

The results so far demonstrated, that motifs both in exon 2 and extended exon 3 mediate the enhancement of AMPA receptors. The conserved tyrosine, which is critically involved in this enhancement, is part of a peptide sequence YEEI, when phosphorylated activates Src family kinases (Lu et al., 1998). When applied to postsynaptic neurons, its activation of Src enhances synaptic transmission (Lu et al., 1998). Furthermore, Src binds to the N-terminus of PSD-95, but not to the other DLG-MAGUKs (Kalia et al., 2006a; Kalia and Salter, 2003). Thus, we hypothesized that differential binding and/or activation of Src family kinases by the N-terminus of the DLG-MAGUKs might cause the differences in AMPA receptor enhancement. To test this hypothesis, we used the Src family kinase activation peptide to test whether its function depends on PSD-95. Notably, this peptide activates Src in *in vitro* assays in absence of any other protein (Kalia et al., 2006b). Thus, the role of PSD-95 in this experiment would be to localize activated Src at the PSD. The peptide (1 mM) was loaded into the patch pipette. 7 min after break in, AMPAR EPSCs of CA1 pyramidal neurons in acute hippocampal slices were recorded (Fig. 4 A-E). In slices from WT mice, the AMPAR EPSCs progressively increased and plateaued after 5-10 min (between group, F_(1,21)_ = 27.39, p < 0.01; group x time, F_(10, 180)_ = 4.17, p < 0.01; Fig. 4B). In simultaneously recorded CA1 pyramidal neurons without peptide, the AMPAR EPSC amplitudes did not change significantly.

**Figure 4:**
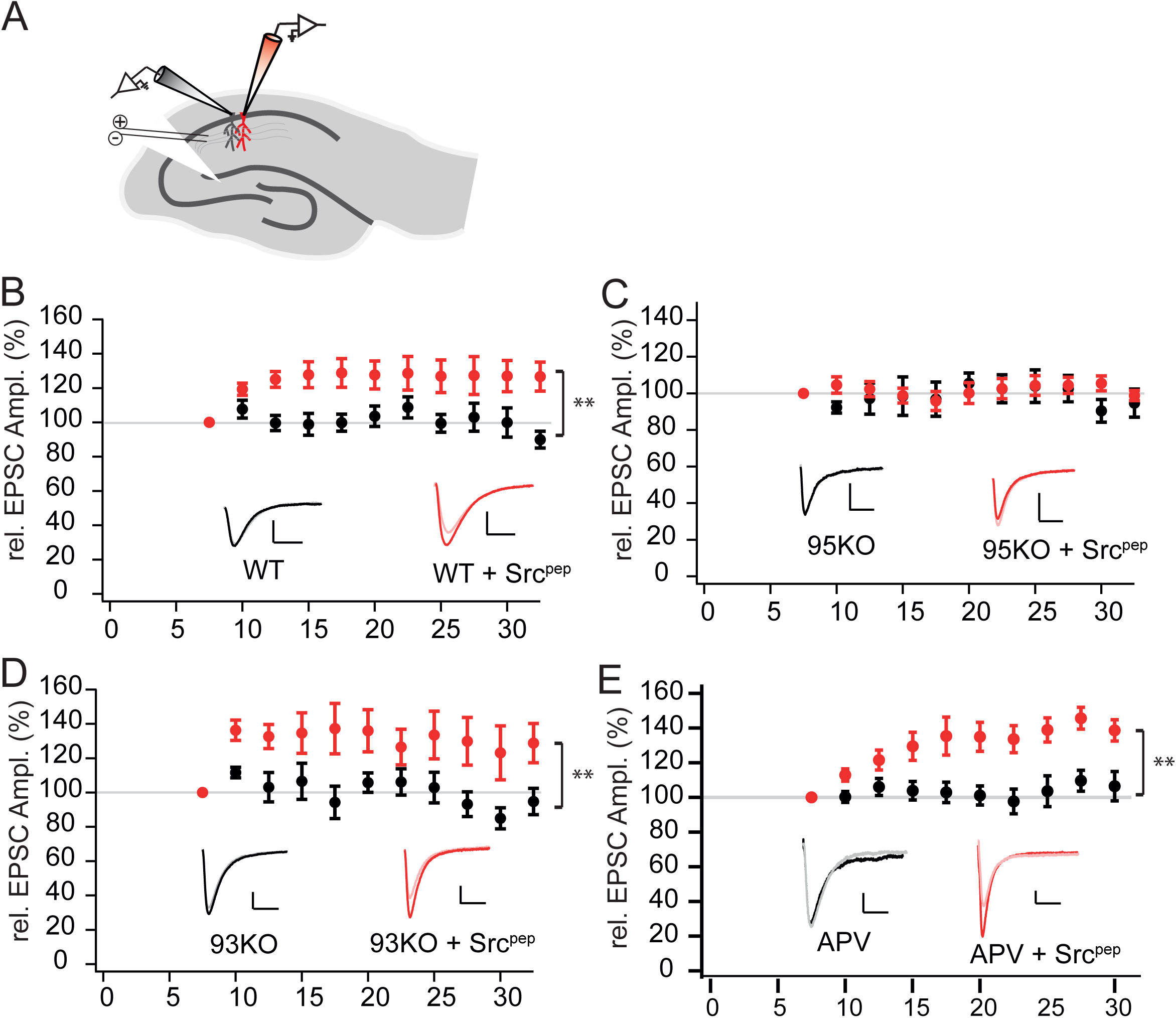
Enhancing function of Src on AMPAR EPSCs requires PSD-95. (A) Schematic drawing of the recording configuration in mouse hippocampal acute slices with control CA1 pyramidal neuron (black) and neighboring CA1 pyramidal neuron with Src activation peptide in intracellular solution (red). (B-E) Normalized AMPAR EPSC in control (black) neuron (B) WT, (C) PSD-95 KO, (D) PSD-93 KO and (E) with 50 µM APV in recording ACSF, and the corresponding neuron with SFK activation peptide (red). Inset shows the sample trace of control neuron EPSC at 7.5 minute (grey) and 30 minute (black), with Src activated neuron EPSC at 7.5 minute (pink) and 30 minute (red). mixed model two-way ANOVA: ** p < 0.01.

In neurons of PSD-95 KO mice, Src activation peptide loading did not enhance AMPAR EPSCs, the magnitude of AMPAR EPSCs was similar to the simultaneously recorded reference neuron (between group, F_(1,20)_ = 0.38, p = 0.54; group x time, F_(9, 176)_ = 1.50, p = 0.15; Fig. 4C), indicating that the Src family kinase enhancing effect was dependent on PSD-95. In contrast, in neurons of PSD-93 KO mice, AMPAR EPSCs were enhanced similarly by the Src family kinase activation peptide as in WT neurons (between group, F_(1,30)_ = 22.14, p < 0.01; group x time, F_(9, 240)_ = 2.85, p < 0.01; Fig. 4D). The most parsimonious explanation of the results is a selective scaffolding of Src by PSD-95 and not PSD-93, consistent with the selective interaction of Src with PSD-95, but not with PSD-93 (Kalia and Salter, 2003).

It was previously reported that enhancing AMPAR EPSCs is mediated by an increase in NMDA receptor function and by lowering the threshold for LTP (Lu et al., 1998). To test whether NMDA receptor activation is required for the enhancing effect of the Src family kinase activation peptide, we repeated the experiment in WT slices in the presence of the NMDA receptor antagonist D-AP5 (between group, F_(1,30)_ = 15.16, p < 0.01; group x time, F_(9, 226)_ = 4.28, p < 0.01; Fig. 4E). Blocking NMDA receptors did not block the effect of the activation peptide on enhancing AMPAR EPSCs, and had also no significant effect on the AMPAR EPSCs of the reference neurons.

We conclude that the Src family kinase activation peptide enhances AMPAR EPSCs. This enhancement requires PSD-95 and is independent of NMDA receptor signaling.

### PSD-95 requires Src to enhance synaptic AMPA receptors

We next tested the opposite, whether the PSD-95α-mediated enhancing effect on AMPAR EPSCs requires Src by comparing PSD-95α over-expression with or without simultaneous knock-down of Src. To generate a loss-of function of Src, we generated a lentivirus with an shRNA to knock-down Src expression that reduced Src efficiently in rat hippocampus primary cultures (Fig. 5A). We over-expressed PSD-95α without shSrc in CA1 pyramidal neurons, and recorded AMPAR EPSCs in parallel from a transduced and neighboring non-transduced neuron (Fig. 5B). Over-expression of PSD-95α enhanced AMPAR EPSCs (n = 14; control, −63.69 ± 8.80 pA; infected, −120.32 ± 14.97 pA, p < 0.01; Fig. 5C, G).

**Figure 5:**
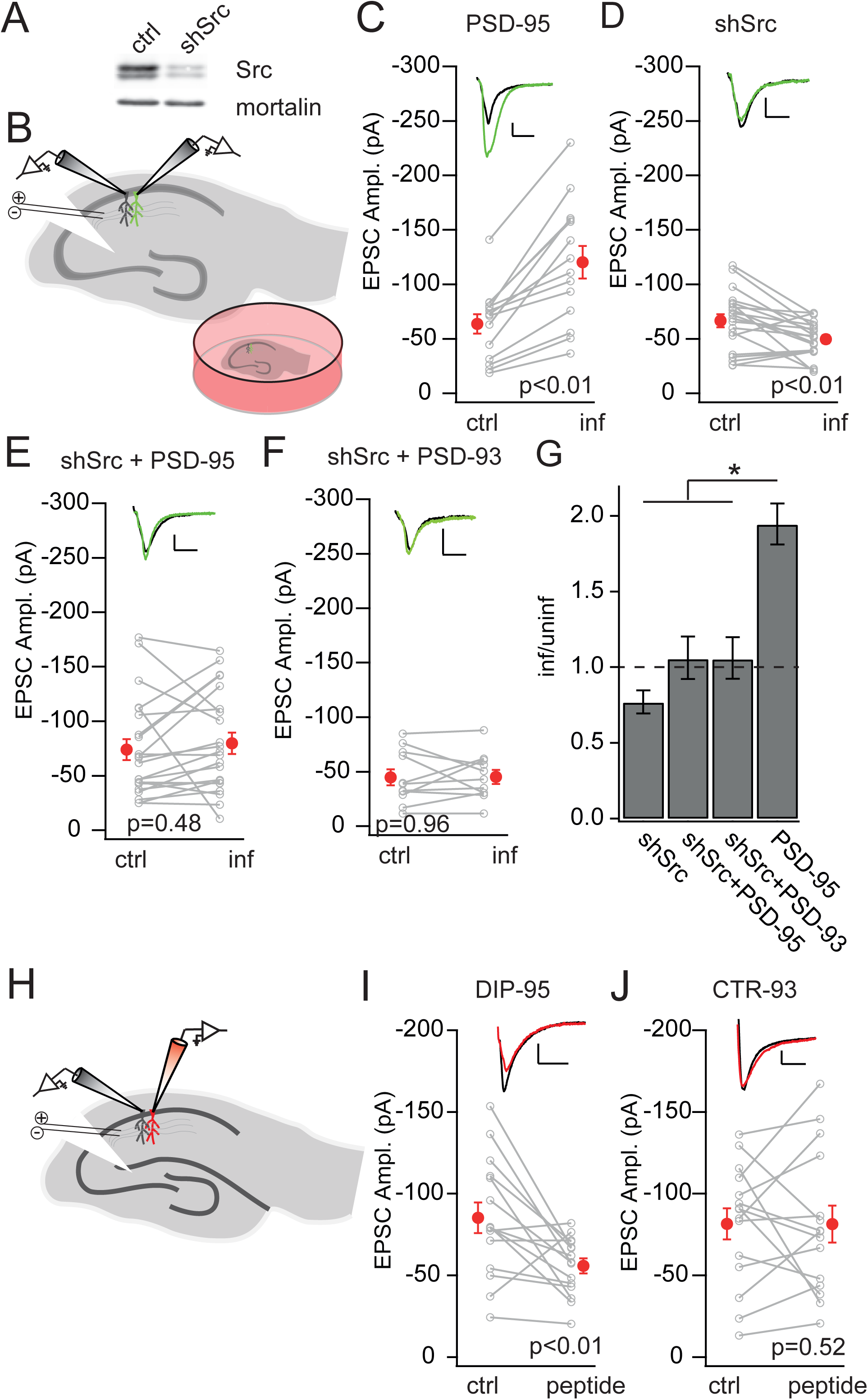
Loss-of Src or peptide disrupting PSD-95/Src interaction inhibit AMPAR EPSCs. (A) Western blot of cell lysates of rat primary hippocampal neuron culture with knockdown of Src versus GFP control neurons. (B) Schematic drawing of the recording configuration of organotypic slices from rat hippocampus. (C-F) Amplitude of AMPAR EPSCs of CA1 pyramidal neurons expressing eGFP-tagged (C) PSD-95α, (D) shRNA against Src, (E) shSrc with PSD-95α over-expression, and (F) shSrc with PSD-93α2 over-expression as lentiviral constructs are plotted against those of simultaneously recorded uninfected neighboring CA1 pyramidal neurons. (G) Summary graph of the comparison between indicated experimental groups of AMPAR EPSC amplitude ratios between transduced (inf) and control neuron (uninf). ANOVA test: * p < 0.05. (H) Schematic drawing of the recording configuration in acute slices of mouse hippocampus with control CA1 neuron (black) and neuron with peptide in the intracellular solution (red). (I-J) Amplitude of AMPAR EPSCs of CA1 pyramidal neurons with (I) dominant interfering peptide of PSD-95 and (J) corresponding PSD-93 peptide in the intracellular solution are plotted against those of simultaneously recorded neighboring control CA1 pyramidal neurons. Insets show sample trace of control neuron (black) and neuron with peptide (red).

Similarly, we expressed the shSrc. AMPAR EPSCs were reduced in shSrc neurons, compared to non-transduced control neurons (n = 20; control, −66.60 ± 6.07 pA; infected, −49.65 ± 3.74 pA, p < 0.01; Fig. 5D, G), indicating that basal synaptic strength requires Src. To test the interaction of Src and PSD-95, we combined the expression of shSrc and PSD-95α in a bi-cistronic lentiviral vector (Schlüter et al., 2006). AMPAR EPSCs of transduced neurons were not significantly different from those of neighboring non-transduced control neurons (n = 22; control, −74.03 ± 9.60 pA; infected, −79.84 ± 9.74 pA, p = 0.48; Fig. 5E, G). Thus, shSrc prevented the enhancing effect of PSD-95α over-expression.

Notably, the AMPAR EPSCs with the combination of shSrc and PSD-95α over-expression were not similarly reduced below control neuron values as with shSrc alone, an effect that we have previously observed with multiple DLG-MAGUK constructs which did not enhance AMPAR EPSCS by themselves but with knock-down of PSD-95, rescued transmission to control neuron level (Schlüter et al., 2006; Xu et al., 2008). So far it is not clear whether it is an additive effect of shSrc reduction and PSD-95-mediated enhancement, or rather knock-down of Src prevents the further enhancement mediated by PSD-95α, thus indicating causality. To distinguish between these possibilities, we expressed PSD-93α2 together with shSrc. Indeed, despite the lack of PSD-93α2 to itself enhance AMPAR EPSCs, the combination brought synaptic transmission back to the control level (n = 11; control, −44.85 ± 7.41 pA; infected, −45.14 ± 6.36 pA, p = 0.96; Fig. 5F), thus behaving similarly as what we observed in previous studies, likely by an unrelated mechanism. When calculating the ratio of AMPAR EPSCs of transduced versus untransduced reference neurons, the enhancing effect of PSD-95α over-expression was significantly different from the effects of the shSrc with PSD-95α, the shSrc with PSD-93α, or the shSrc alone group, while there was no significantly different between the other groups (PSD-95, 1.94 ± {+0.14, −0.13}; shSrc, 0.77 ± {+0.08, −0.07}; shSrc + PSD-95, 1.05 ± {+0.15, - 0.13}, shSrc + PSD-93, 1.05 ± {+0.15, −0.13}; F_(3,63)_ = 10.33, p < 0.01; PSD-95 vs. shSrc, p < 0.01; PSD-95 vs. shSrc + PSD-95, p < 0.01; shSrc + PSD-95 vs. shSrc + PSD-93, p = 1; shSrc vs. shSrc + PSD-95, p = 0.15; Fig. 5G). These results indicated that PSD-95α lost its enhancing effect on synaptic AMPARs when Src expression was absent, thus behaves similar to PSD-93α2. In conclusion, Src is required for both, maintaining basal synaptic transmission and for the effect of PSD-95α on enhancing AMPAR EPSCs.

### Continuous Src tethering to PSD-95 is required to maintain AMPAR EPSCs

We found so far that the Src family kinase activation peptide enhanced AMPAR EPSCs already after 10 min (Fig. 4). We concluded that the function of PSD-95α is to tether active Src to the PSD to enhance AMPAR EPCs. Furthermore, the relative fast action of the peptide indicated that Src activity is under dynamic control, and might thus be continuously activated to maintain synaptic strength. To test this hypothesis, we used a dominant interfering peptide (DIP) procedure. The PSD-95 DIP (peptide sequence: DTLEAPGYELQVNGT) was previously shown to disrupt the PSD-95/Src interaction (Kalia et al., 2006b). We loaded this peptide (1 mM) into the patch-pipette and recorded AMPAR EPSCs in comparison to a parallel recorded reference cells without the peptide (Fig. 5H). We then averaged the AMPAR EPSCs after at least 20 min break in. The AMPAR EPSCs with the PSD-95 DIP were reduced compared to the reference neurons (n = 15; control, - 85.23 ± 9.38 pA; DIP, −55.89 ± 4.54 pA, p < 0.01; Fig. 5I), indicating that the PSD-95/Src interaction is required to maintain the strength of AMPAR EPSCs.

For control, we used the corresponding sequence of PSD-93α2 as a putative PSD-93 DIP (peptide sequence: DTLDTIPYVNGT). AMPAR EPSCs of neurons loaded with the PSD-93 DIP were not significantly different compared to the simultaneously recorded reference neuron without peptide (n = 15; control, −81.58 ± 9.52 pA; infected, −81.43 ± 11.35 pA, p = 0.99; Fig. 5J). Together these results indicate that the interaction of PSD-95 with Src is required to maintain the function of synaptic AMPA receptors, presumably by tethering the active Src at the PSD to maintain the tyrosine phosphorylation of substrate proteins.

## Discussion

We found that enhancing AMPAR EPSCs depends mutually on PSD-95α and Src. Using domain swapping and comparative analysis of amino acids, we identified a conserved tyrosine in PSD-95α and SAP97α that is a phenylalanine in PSD-93α2, coded by the extended exon 3 that together with additional motifs in the N-terminus mediated the enhancing effect (Fig. 2, 3). The tyrosine if phosphorylated is part of a known consensus Src activation peptide motif that binds to the Src SH2 domain, and is localized immediately downstream of a previously described Src interaction motif in PSD-95α (Fig. 1)(Kalia et al., 2006b; Lu et al., 1998). Blocking the PSD-95/Src interaction with a peptide of the interaction motif during recordings reduced EPSCs, indicating that the continuous tethering of Src by PSD-95 was required to maintain synaptic strength (Fig. 5). Together these results support a model in which Src is tethered to the PSD by PSD-95α in a signaling complex, and the PSD-95 Src activation motif activates Src to maintain synaptic AMPA receptor function.

### Differential function of DLG-MAGUK α-isoforms

Using N-terminal domains swapping between PSD-93α2 and PSD-95α or SAP97α, respectively, we narrowed down motifs to a single amino acid in the N-terminus that mediate part of the enhancing effect of PSD-95α and SAP97α on AMPAR EPSCs. The Y63 or Y93 is conserved in PSD-95α and SAP97α, respectively, while it is the F103 in PSD-93α2. The tyrosine is part of the motif YEEI which was shown, if phosphorylated in a peptide, to bind to the SH2 domain of Src family kinases and to activate them (Lu et al., 1998). Swapping the tyrosine and phenylalanine between PSD-95α and PSD-93α2 transferred the enhancing effect with the tyrosine onto either protein (Fig. 3). The magnitude of the enhancing effect was similar to the swap of the extended exon 3, but smaller than the full N-terminal swap, indicating that the tyrosine is only one part of what mediates the enhancing function.

Upstream of the putative Src activation motif in PSD-95α, is a previously described motif, coded by exon 3 that also binds to the SH2 domain of Src, although non-canonically, as binding does not require a phospho-Tyr (Kalia et al., 2006b). A peptide of this motif decreased AMPAR EPSCs, while the corresponding peptide of PSD-93α2 did not (Fig. 5), a result that is consistent with its role as a dominant interfering peptide to prevent the PSD-95/Src interaction. However, it was previously shown that the PSD-95-derived peptide competes with the Src activation peptide for binding to the Src SH2 domain, thereby inhibits Src activity (Kalia et al., 2006b). But, it is not clear whether the PSD-95-derived peptide interferes with the tethering of Src to native PSD-95α by blocking the interaction domain, thus prevent binding of the activation motif in native PSD-95α to the Src SH2 domain, or rather the two peptides compete for Src SH2 binding. Whatever the scenario, our results indicate that the tyrosine of the YEEI motif enhances AMPAR EPSCs and the effect of the PSD-95 DIP supports a tethering function of the upstream motif.

Sequence comparisons of coded amino acids of exon 2 did not reveal a similar candidate as the tyrosine, coded by the extended exon 3 (Fig. 1). Thus, the function of exon 2 remains speculative. Previous studies reported that phosphorylation by the cyclin-dependent kinase 5 (cdk5) of PSD-95 reduces its interaction with Src (Zhang et al., 2008). Cdk5 is predicted to phosphorylate T19, S25, and S35 of PSD-95α (Morabito et al., 2004). However, the N-terminal cdk5 phosphorylation sites differ among PSD-95, SAP97 and PSD-93 (Fig. 1). The T19 is unique to PSD-95α, while PSD-95α S25 is conserved in SAP97α, and conversely PSD-95α S35 is conserved in PSD-93α2. Thus, there is no clear pattern of conservation between PSD-95α and SAP97α. Nevertheless, it was reported that the phosphorylation of PSD-95α T19 promotes the interaction of the N-terminus with Ca^2+^/calmodulin (Zhang et al., 2014). T19 might thus uniquely regulate PSD-95α synaptic localization by Ca^2+^/calmodulin. For Src activation, the second phosphorylation site which is conserved between PSD-95α and SAP97α might be critical. Future studies will need to clarify whether these sites constitute the other part for PSD-95α to enhance AMPAR EPSCs.

While the N-terminal swaps narrowed down the motifs in PSD-95α and SAP97α that mediate the enhancement of AMPAR EPSCs and let us identify the role of Src in this process, a number of observations indicate that the C-terminal domain of PSD-93α2 might have an inhibitory role in AMPAR function. Either motif coded by *Dlg2* exon 2 or extended exon 3 if swapped into SAP97α did not reduce the enhancing effect of mutant SAP97α (Fig. 2), indicating that the remaining N-terminal motifs in SAP97α were sufficient to enhance AMPAR EPSCs and that neither of the PSD-93 motifs contained an active inhibitory function. The sufficiency of one of the motifs for enhancement is reminiscent of PSD-95α that lacked amino acids 46 to 64, thus most of the Src interaction and activation motif of the extended exon 3, but was still capable to enhance AMPAR EPSCs (Xu et al., 2008). Thus, if the C-terminal domain is from PSD-95α or SAP97α, one of the N-terminal halves is sufficient for enhancing AMPAR EPSCs. In contrast, if the C-terminal part is from PSD-93, N-terminal motifs coded by both *Dlg1* exon 2 or extended exon 3 were required for the full enhancing effect (Fig. 2). Furthermore, if the N-terminal domain of PSD-95α was swapped onto the C-terminal part of PSD-93, then the magnitude of enhancement was smaller as with PSD-95α over-expression (Fig. 1). We interpret this result as evidence that the C-terminal part of PSD-93 contains an active inhibitory function for synaptic AMPA receptors. Indeed, we observed that over-expression of PSD-93α2 reduced synaptic AMPA receptor function in the visual cortex but less consistently in the hippocampus (Favaro et al., 2018; Krüger et al., 2013).

### Role of Src in enhancing synaptic transmission

We found that synaptic AMPAR EPSCs depend mutually on Src and PSD-95α (Fig. 4, 5). This conclusion is supported by the following results. First, the enhancing effect of the Src family kinase activation peptide was prevented in PSD-95 KO mice, but not PSD-93 KO mice (Fig. 4). This peptide mediates the enhancing effect via Src activation, as a Src-specific inhibitory peptide blocks this enhancement (Lu et al., 1998). Conversely, the enhancing effect of PSD-95α over-expression was prevented by knock-down of Src (Fig. 5). Second, the Src family kinase activation peptide enhanced AMPAR EPSCs independent of NMDA receptor activation (Fig. 4), thus similar to the PSD-95α enhancing effect on AMPAR EPSCs, does not depend on NMDA receptor activation (Schlüter et al., 2006; Zhang and Lisman, 2012). Third, a peptide mimicking the PSD-95α binding site to Src reduced AMPAR EPSCs (Fig. 5). Together these results support a model in which PSD-95α tethers Src to the PSD and promotes its activation potentially via the phosphorylated YEEI Src SH2 binding motif.

A previous study reported that the enhancement of AMPAR EPSCs is mediated by Src’s effect on NMDA receptors (Lu et al., 1998). The open channel blocker MK-801 blocked the enhancing effect of the phosphorylated YEEI peptide. It is not mutually exclusive that Src enhances AMPAR EPSC by facilitating NMDA receptor signaling as well as through NMDA receptor independent mechanisms. Despite that, the action of MK-801 and D-AP5 on NMDA receptors differ. It has been shown that MK-801 blocks ionotropic signaling of NMDA receptors, but not metabotropic NMDA receptor signaling, while D-AP5 blocks both (Carter and Jahr, 2016; Nabavi et al., 2013; Stein et al., 2015). Thus, Src activation for NMDA-dependent and independent enhancement of AMPA receptor function might follow different signaling pathways.

In further support of this notion, the mechanism of Src-mediated AMPA receptor enhancement appears different to the better known role of Src to regulate NMDA receptor activity during long-term synaptic potentiation (Salter and Kalia, 2004). mGluR signaling is involved in the enhancing effect of PSD-95α over-expression on AMPAR EPSCs and GluN2A/2B phosphorylation (Heidinger et al., 2002; Zhang and Lisman, 2012). However, the signaling pathways differ. mGluR1 signaling leads to Src activation and GluN2A and 2B phosphorylation, while mGluR5 signaling was reported to be required for the enhancing function of PSD-95α on AMPAR EPSCs. G-protein-coupled receptor activation has been linked to Src activation and mGluR5 signaling to Src family kinase activation (Benquet et al., 2002; Lu et al., 1999), but whether mGluR5 signals via Src to enhance AMPAR EPSCs remains elusive.

The interpretation is further complicated by the report that PSD-95/Src interaction has a negative role on NMDA receptor phosphorylation (Kalia et al., 2006b), while another study reports that PSD-95/Src binding enhances NMDA receptor phosphorylation and synaptic expression (Zhang et al., 2008). Furthermore, contrary to this inhibitory role of PSD-95, over-expression of PSD-95α has only a minor and a rather enhancing effect on NMDAR EPSCs (Elias et al., 2006; Nakagawa et al., 2004; Schlüter et al., 2006; Schnell et al., 2002a). Thus further studies will have to identify the mechanistic differences in synaptic regulation of AMPAR and NMDAR EPSCs by Src. In conclusion, a number of signaling pathways have been identified to activate Src and enhance synaptic NMDA and AMPA receptors, whereas our results identify motifs in the N-terminus of PSD-95α to be essential for Src to enhance and maintain synaptic AMPAR function. This results support the role of PSD-95 as signaling scaffold to regulate AMPA receptor synaptic incorporation.

## Acknowledgement

We thank S. Ott-Gebauer for excellent technical assistance, the AGCT core facility for primer synthesis and DNA sequencing. This work was funded by the National Institute for Neurological Disorders and Stroke (NIH NS107604), and the Whitehall Foundation 2018-05-68 to O.M.S. The European Neuroscience Institute Göttingen is jointly funded by the Göttingen University Medical School and the Max Planck Society.

